# Neutralization of N501Y mutant SARS-CoV-2 by BNT162b2 vaccine-elicited sera

**DOI:** 10.1101/2021.01.07.425740

**Authors:** Xuping Xie, Jing Zou, Camila R. Fontes-Garfias, Hongjie Xia, Kena A. Swanson, Mark Cutler, David Cooper, Vineet D. Menachery, Scott Weaver, Philip R. Dormitzer, Pei-Yong Shi

## Abstract

Rapidly spreading variants of SARS-CoV-2 that have arisen in the United Kingdom and South Africa share the spike N501Y substitution, which is of particular concern because it is located in the viral receptor binding site for cell entry and increases binding to the receptor (angiotensin converting enzyme 2). We generated isogenic N501 and Y501 SARS-CoV-2. Sera of 20 participants in a previously reported trial of the mRNA-based COVID-19 vaccine BNT162b2 had equivalent neutralizing titers to the N501 and Y501 viruses.

## Main text

We previously reported that BNT162b2, a nucleoside modified RNA vaccine that encodes the SARS-CoV-2 full length, prefusion stabilized spike glycoprotein (S), elicited dose-dependent SARS-CoV-2–neutralizing geometric mean titers (GMTs) that were similar to or higher than the GMT of a panel of SARS-CoV-2 convalescent human serum samples.^1^ We subsequently reported that, in a randomized, placebo-controlled trial in approximately 44,000 participants 16 years of age or older, a two-dose regimen of BNT162b2 conferred 95% protection against Covid-19.^2^

Since the previously reported studies were conducted, rapidly spreading variants of SARS-CoV-2 have arisen in the United Kingdom and South Africa.^3,4^ These variants have multiple mutations in their S glycoproteins, which are key targets of virus neutralizing antibodies. These rapidly spreading variants share the spike N501Y substitution. This mutation is of particular concern because it is located in the viral receptor binding site for cell entry, increases binding to the receptor (angiotensin converting enzyme 2), and enables the virus to expand its host range to infect mice.^5,6^

We generated an isogenic Y501 SARS-CoV-2 on the genetic background of the N501 clinical strain USA-WA1/2020, which also provided the genetic background of the BNT162b2-encoded spike antigen. Sera of 20 participants in the previously reported trial,^1,2^ drawn 2 or 4 weeks after immunization with two 30-μg doses of BNT162b2 spaced three weeks apart, were tested for neutralization of N501 and Y501 viruses by a 50% plaque reduction neutralization assay (PRNT50; Figure 1). The ratio of the 50% neutralization GMT of the sera against the Y501 virus to that against the N501 virus was 1.46, indicating no reduction in neutralization activity against the virus bearing the Y501 spike (Supplementary material).

**Figure 1.**
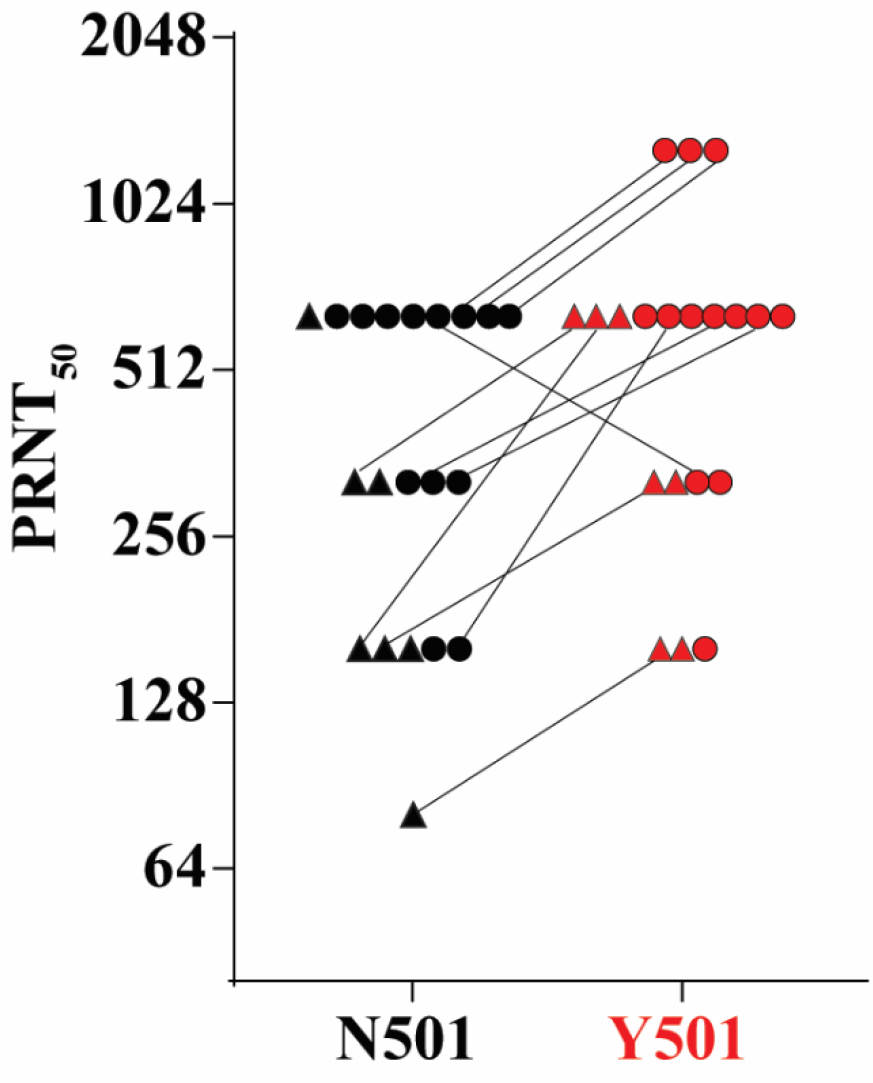
50% plaque reduction neutralization titers of 20 sera from BNT162b2 vaccine recipients against N501 and Y501 SARS-CoV-2. Seven sera (indicated by triangles) were drawn 2 weeks after the second dose of vaccine; 13 sera (indicated by circles) were drawn 4 weeks after the second dose.

A limitation of this finding is that the Y501 virus does not include the full set of spike mutations found on the rapidly spreading strains in the UK or South Africa.^3,4^ Nevertheless, preserved neutralization of Y501 virus by BNT162b2-elicited human sera is consistent with preserved neutralization of a panel of 15 pseudoviruses bearing spikes with other mutations found in circulating SARS-CoV-2 strains.^7^

The ongoing evolution of SARS-CoV-2 necessitates continuous monitoring of the significance of changes for vaccine coverage. This surveillance is accompanied by preparations for the possibility that a future mutation in SARS-CoV-2 might necessitate a vaccine strain change. Such a vaccine update would be facilitated by the flexibility of mRNA-based vaccine technology.

## Acknowledgments

Supported by Pfizer and BioNTech. We thank colleagues at Pfizer, BioNTech, and UTMB for helpful discussions and support during the study. We thank the Pfizer-BioNTech clinical trial C4591001 participants, from whom the post-immunization human sera were obtained. We thank the many colleagues at Pfizer and BioNTech who developed and produced the BNT162b2 vaccine candidate. P.-Y.S. was supported by NIH grants AI142759, AI134907, AI145617, and UL1TR001439, and awards from the Sealy & Smith Foundation, Kleberg Foundation, the John S. Dunn Foundation, the Amon G. Carter Foundation, the Gilson Longenbaugh Foundation, and the Summerfield Robert Foundation.

## Data availability

The data that support the findings of this study are available from the corresponding authors upon reasonable request.

## Author contributions

Conceptualization, X.P., V.D.M., S.W., P.-Y.S.; Methodology, X.P., J.Z., C.R.F.G., H.X.,

P.-Y.S; Investigation, X.P., J.Z., C.R.F.G., H.X., K.A.S., D.C., P.R.D., P.-Y.S; Resources, M.C., D.C., P.R.D., P.-Y.S; Data Curation, X.P., J.Z., C.R.F.G., P.-Y.S; Writing-Original Draft, X.P., P.-Y.S; Writing-Review & Editing, X.P., P.R.D., P.-Y.S.; Supervision, X.P., M.C., D.C., P.R.D., P.-Y.S.; Funding Acquisition P.-Y.S.

## Competing financial interests

X.X., V.D.M., and P.-Y.S. have filed a patent on the reverse genetic system. K.A.S., M.C., D.C., and P.R.D. are employees of Pfizer and may hold stock options. X.P., J.Z., C.R.F.G., H.X., and P.-Y.S. received compensation from Pfizer to perform the neutralization assay.

## References

1. Walsh EE, Frenck RW, Jr., Falsey AR, et al. Safety and Immunogenicity of Two RNA-Based Covid-19 Vaccine Candidates. N Engl J Med 2020.

2. Polack FP, Thomas SJ, Kitchin N et al. Safety and efficacy of the BNT162b2 mRNA Covid-19 vaccine. N Eng. J Med 2020. DOI: 10.1056/NEJMoa2034577.

3. Volz E, Mishra S, Chand M et al. Report 42 - Transmission of SARS-CoV-2 Lineage B.1.1.7 in England: Insights from linking epidemiological and genetic data. https://www.imperial.ac.uk/mrc-global-infectious-disease-analysis/covid-19/report-42-sars-cov-2-variant/.

4. Tegally H, Wilkinson E, Giovanetti M et al. Emergence and rapid spread of a new severe acute respiratory syndrome-related coronavirus 2 (SARS-CoV-2) lineage with multiple spike mutations in South Afric. medRxiv 2020. https://doi.org/10.1101/2020.12.21.20248640

5. Gu H, Chen Q, Yang G, et al. Adaptation of SARS-CoV-2 in BALB/c mice for testing vaccine efficacy. Science 2020;369:1603–7.

6. Chan KK, Tan TJC, Narayanan KK & Procko E. An engineered decoy receptor for SARS-CoV-2 broadly binds protein S sequence variants. Cold Spring Harbor Laboratory 2020.doi: 10.1101/2020.10.18.344622

7. Sahin U, Muik A, Volger I et al., BNT162b2 induces SARS-CoV-2 -neutralising antibodies and T cells in humans. medRxiv 2020. doi: https://doi.org/10.1101/2020.12.09.20245175

