## Supplementary material for "Neutralization of N501Y mutant SARS-CoV-2 by BNT162b2 vaccine-elicited sera"

### Table of Contents


**Materials and Methods**

- Construction of isogenic viruses
- Neutralization assay

**Figures**

- Figure S1. Diagram of the N501Y substitution.
- Figure S2. Plaque morphologies of N501 and Y501 viruses.
- Figure S3. Scheme of the BNT162 vaccination and serum sampling.
- Figure S4. Plot of the ratio of $PRNT_{50}$ between Y501 and N501 viruses.

**Tables**

- Table S1. $PRNT_{50}$ values of 20 BNT162b2 post-immunization sera against N501 and Y501 SARS-CoV-2.

**References**

### Materials and Methods

#### Construction of isogenic viruses

We prepared an isogenic pair of SARS-CoV-2 containing the N501 or Y501 spike protein (Figure S1). The N501Y mutation was generated by an A-to-T substitution at nucleotide 23,063 of the viral genome using an infectious cDNA clone of clinical strain WA1 (2019-nCoV/USA_WA1/2020).[1] Following a previously reported mutagenesis protocol,[2] we recovered N501 and Y501 viruses with titers of >$10^7$ plaque-forming units (PFU) per ml. The two viruses developed similar plaque morphologies on Vero E6 cells (Fig. S2).

#### Serum specimens and neutralization assay

The immunization and serum collection regimen is illustrated schematically in Fig. S3. For measuring neutralization titers, each serum was 2-fold serially diluted in culture medium with the first dilution of 1:40 (dilution range of 1:40 to 1:1280). The diluted serum was incubated with 100 PFU of N501 or Y501 virus at 37 °C for 1 h, after which the serum-virus mixtures were inoculated onto Vero E6 cell monolayer in 6-well plates. A conventional (non-fluorescent) plaque reduction neutralization assay was performed to quantify the serum-mediated virus suppression as previously reported.[3] A minimal serum dilution that suppressed >50% of viral plaques is defined as $PRNT_{50}$. A table of the neutralization titers is provided (Table S1). The ratio for each serum of the $PRNT_{50}$ against N501 and Y501 virus is plotted in Fig. S4.

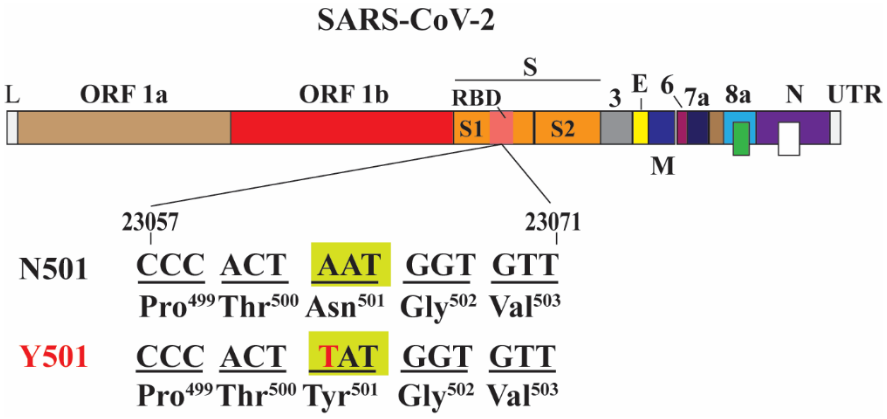


**Figure S1.** Diagram of the N501Y substitution. L – leader sequence; ORF – open reading frame; RBD – receptor binding domain; S – spike glycoprotein; S1 – N-terminal furin cleavage fragment of S; S2 – C-terminal furin cleavage fragment of S; E – envelope protein; M – membrane protein; N – nucleoprotein; UTR – untranslated region.

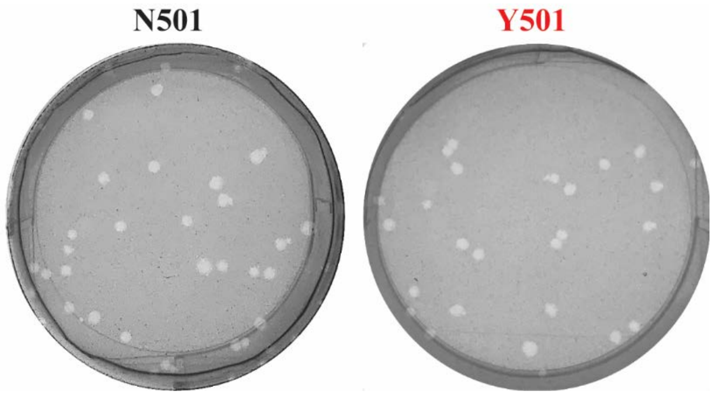


**Figure S2.** Plaque morphologies of N501 and Y501 SARS-CoV-2 on Vero E6 cells.

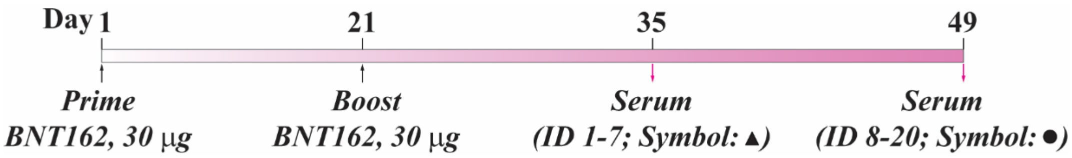


**Figure S3.** Scheme of the BNT162 vaccination and serum sampling.

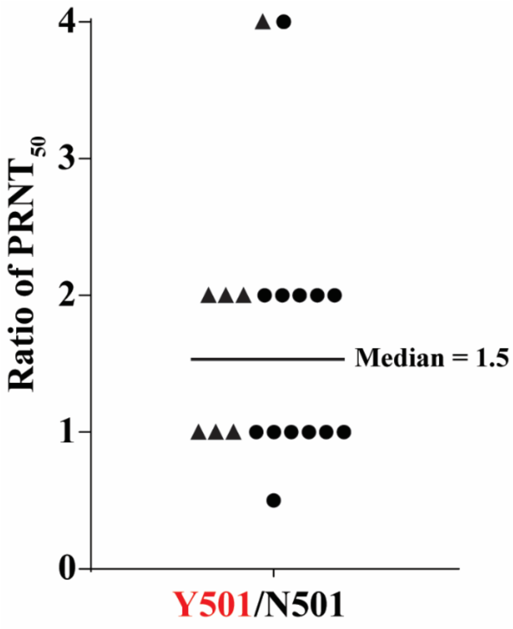


**Figure S4.** Plot of the ratio of $PRNT_{50}$ between Y501 and N501 viruses. Triangles represent sera drawn two weeks after the second dose; circles represent sera drawn four weeks after the second dose.

**Table S1**. $PRNT_{50}$ values of 20 BNT162b2 post-immunization sera against N501 and Y501 SARS-CoV-2.

| Serum ID | $PRNT_{50}$ | | $PRNT_{50}$ ratio (Y501/N501) |
|---|---|---|---|
| | N501 | Y501 | |
| 1 | 160 | 640 | 4 |
| 2 | 160 | 320 | 2 |
| 3 | 320 | 640 | 2 |
| 4 | 80 | 160 | 2 |
| 5 | 160 | 160 | 1 |
| 6 | 320 | 320 | 1 |
| 7 | 640 | 640 | 1 |
| 8 | 160 | 160 | 1 |
| 9 | 640 | 640 | 1 |
| 10 | 640 | 1280 | 2 |
| 11 | 160 | 640 | 4 |
| 12 | 320 | 320 | 1 |
| 13 | 640 | 1280 | 2 |
| 14 | 640 | 320 | 0.5 |
| 15 | 320 | 640 | 2 |
| 16 | 320 | 640 | 2 |
| 17 | 640 | 640 | 1 |
| 18 | 640 | 1280 | 2 |
| 19 | 640 | 640 | 1 |
| 20 | 640 | 640 | 1 |

### Supplementary References

1. Xie X, Muruato A, Lokugamage KG, et al. An Infectious cDNA Clone of SARS-CoV-2. Cell Host Microbe 2020;27:841-8 e3.
2. Plante JA, Liu Y, Liu J, et al. Spike mutation D614G alters SARS-CoV-2 fitness. Nature 2020.
3. Muruato AE, Fontes-Garfias CR, Ren P, et al. A high-throughput neutralizing antibody assay for COVID-19 diagnosis and vaccine evaluation. Nat Commun 2020;11:4059.
